# ImSpiRE: Image feature-aided spatial resolution enhancement method

**DOI:** 10.1101/2023.05.04.539342

**Authors:** Yuwei Hua, Yizhi Zhang, Zhenming Guo, Shan Bian, Yong Zhang

## Abstract

The resolution of most spatially resolved transcriptomic technologies usually cannot attain the single-cell level, limiting their applications in biological discoveries. Here, we introduce ImSpiRE, an image feature-aided spatial resolution enhancement method for *in situ* capturing spatial transcriptome. Taking the information stored in histological images, ImSpiRE solves an optimal transport problem to redistribute the expression profiles of spots to construct new transcriptional profiles with enhanced resolution, together with imputing the gene expression profiles in unmeasured regions. Applications to multiple datasets confirm that ImSpiRE can enhance spatial resolution to the subspot level while contributing to the discovery of tissue domains, signaling communication patterns, and spatiotemporal characterization.

## Main

Multiple cells of various types are structurally organized within the complex spatial architecture of tissues to perform biological functions. The acquisition of the spatial transcriptome in tissues is essential to reveal spatially specific gene expression and structural patterns. Over the past decade, various forms of *in situ* hybridization (ISH) technologies have been used to directly visualize the spatial context of mRNA transcripts in fluorescence images, such as seqFISH[1, 2], seqFISH+[3] and MERFISH[4]. These approaches can be practically applied to only a small field of view while having low throughput and complex custom protocols, thus limiting their widespread applications[5]. In contrast, because of mature commercial platforms, *in situ* capturing (ISC) technologies, which capture transcripts *in situ* and then perform *ex situ* sequencing by using specific spatial barcodes, have been the most widely promoted[6]. The strategy of ISC allows for an unbiased analysis of the transcriptome in tissue sections[7]. By using various spatial barcodes, ISCs measure the gene expression levels at measurement locations known as spots. Spatial transcriptomics (ST), for instance, contains ∼1,000 spots in a microarray[8]. A spot in ST is 100 μm in diameter and typically composed of 10-40 cells[7]. Visium Spatial Gene Expression (Visium) from 10x Genomics increases the number of spots to ∼5,000 and reduces the diameter size to 55 μm per spot, but the resolution still cannot achieve the single-cell level. Stereo-seq[9] improves the size of measurement locations to below 1 μm. However, its limited RNA capture efficiency causes multiple measurement locations to be combined into a single spot for analysis, which largely reduces the spatial resolution. Therefore, the spatial transcriptome derived from existing ISC spatial transcriptome techniques measures the total gene expression of the cell mixture at each spot, and such a mixture is a barrier to the investigation of complex tissue architecture.

To reduce the effects of mixed measurements caused by low spatial resolution, multiple computational methods have been developed. A common strategy is cell-type deconvolution within spots, such as CARD[10], STRIDE[11], RCTD[12], SPOTlight[13], etc. Given the complementary information provided by the ISC spatial transcriptome and scRNA-seq, the proportions of cell types within a spot can be inferred. The major drawback of this strategy is the reliance on additional single-cell transcriptomes and a priori knowledge of cell types. In addition, the inferred cell type proportions do not directly improve spatial resolution. Different from deconvolution methods, BayesSpace[5] subdivides a spot into multiple equal-sized subspots and infers the gene expression at the subspot level from its spatial neighborhoods without using external information. However, the resolution enhancement of BayesSpace is limited to the density of original spots, and the information from spatial neighborhoods is uncorrelated when their distances are far. Two recent methods, XFuse[14] and DeepSpaCE[15], fuse spatial transcriptome and parallel histology images of tissues to impute gene expression profiles in regions not measured by spots. However, neither method enhances the spatial resolution, regardless of whether the regions are measured. To the best of our knowledge, there has been no computational method that can enhance spatial resolution while imputing the gene expression profiles for unmeasured regions in ISC spatial transcriptomes.

Here, we propose ImSpiRE, an image feature-aided spatial resolution enhancement method for ISC spatial transcriptomes. ImSpiRE first defines patches that are densely arranged on the measured and unmeasured regions of the tissue section. Then, ImSpiRE extracts image features and coordinates from spots and patches in the parallel histology image. ImSpiRE calculates an optimal probabilistic embedding that redistributes the expression profiles of spots to the patches to construct the transcriptional profiles with enhanced resolution. ImSpiRE does not require additional single-cell data or prior knowledge, and it can impute the expression profiles on the regions not measured by original spots. Through applications to several published ISC spatial transcriptomes, we demonstrate that ImSpiRE can effectively enhance the spatial resolution of the transcriptome to the subspot level, which can aid in revealing the complex spatial architecture of biological tissues.

## Results

### Nearby spots with similar image features show similar transcriptional profiles

To enhance the spatial resolution of ISC spatial transcriptomes, we propose the following assumptions. First, nearby spots share similar transcriptional profiles within a tissue section. The assumption is based on the fact that nearby cells tend to share similar abundances of nutrients and signaling molecules[16]. Second, spots with consistent image features in the parallel histological image tend to have similar transcriptional profiles. The assumption is based on the intuitive idea that cells with similar transcriptional profiles tend to share similar visual phenotypes (such as cell morphology, density and staining intensity). To evaluate the above assumptions, we used a public Visium dataset on a mouse brain sagittal posterior section, which contains 3,355 spots and a parallel H&E-stained image; the spots were clustered into 11 clusters (Fig. 1A). For the first assumption, we calculate the physical distances and expression distances between all spots by their spatial coordinates and transcriptional profiles, respectively. We observed that all spots in the same cluster tended to be located close to each other in the tissue section (Fig. 1B). We further found that the average expression distances between spots gradually increased with increasing physical distances (Fig. 1C). Furthermore, the Pearson correlation coefficients (PCCs) between the expression distances and the physical distances for all spot pairs were significantly lower than those for random matching (Fig. 1D). For the second assumption, we cropped the fixed size subimages for all spots according to their coordinates in the parallel H&E-stained image and extracted the image features (such as intensity and texture) for each spot (Fig. 1E). By calculating the image feature distances, we observed that spots in the same clusters tended to show similar image features (Fig. 1F). Moreover, there is a clear trend of increasing expression distances with increasing image feature distances (Fig. 1G). The PCCs between the expression distances and the image feature distances are significantly higher than those of random matching (Fig. 1H). We evaluated multiple ISC spatial transcriptomes and observed consistent results (Fig. S1). Our results confirm the assumptions in multiple ISC spatial transcriptomes that nearby spots with similar image features tend to have similar transcriptional profiles.

**Fig. 1.**
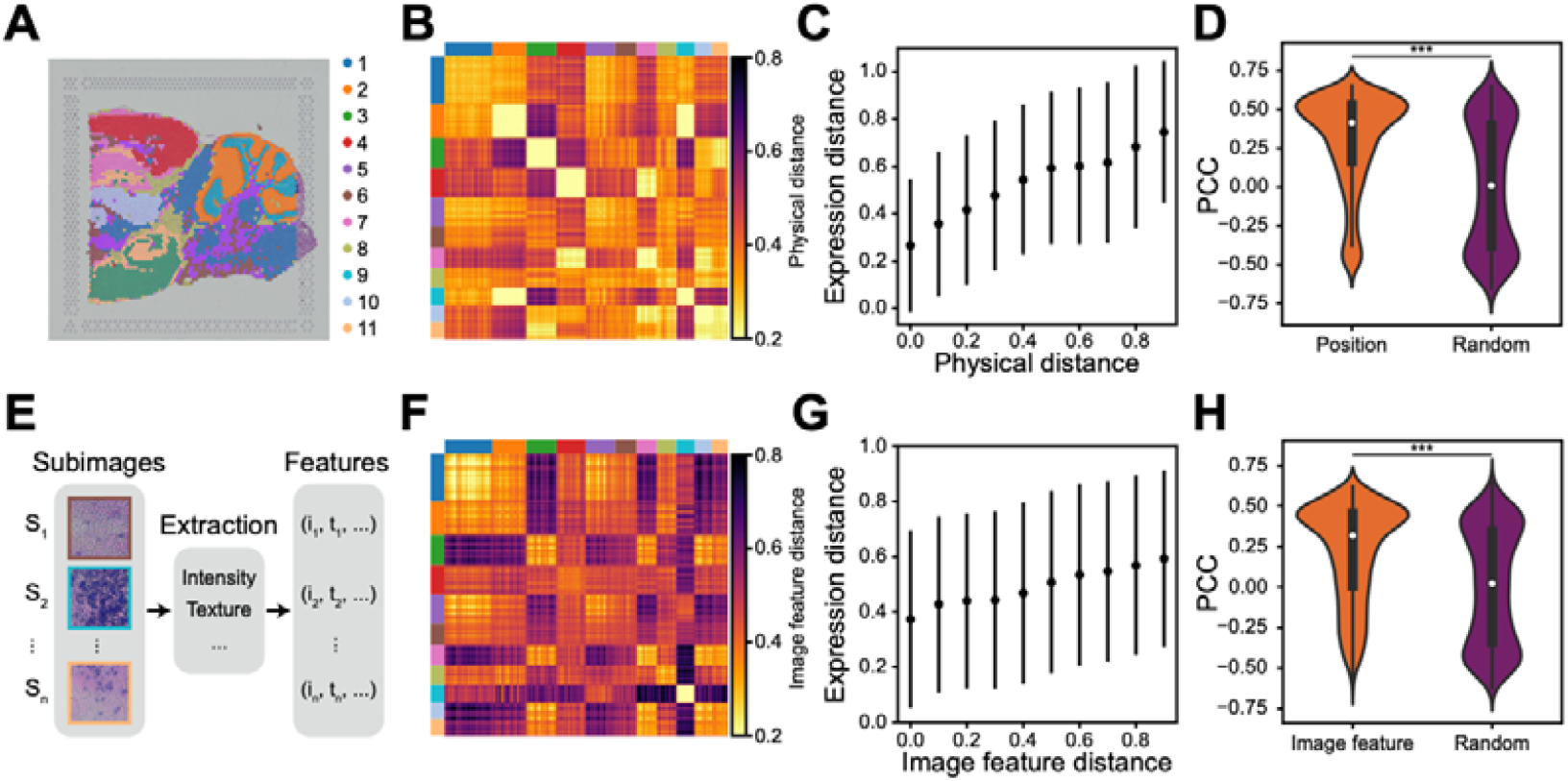
Nearby spots with similar image features show similar transcriptional profiles. **A**. The coordinates of spots on parallel H&E-stained image of mouse brain sagittal posterior section. The spots are clustered into eleven clusters in the original study. The eleven colors represent different clusters. **B**. Heatmap shows that all spots in the same cluster tend to be located close to each other in the tissue section. The physical distances are normalized Euclidean distances of spatial coordinates between all spot pairs. The colors are consistent with eleven clusters of spots in A. **C**. The plot shows the relationship between the expression distances and physical distances for spot pairs. The expression distances are normalized correlation-based distances of transcriptional profiles between all spot pairs. The values in the plot are average expression distances s.d. **D**. The Pearson correlation coefficients (PCCs) between expression distances and physical distances for all spot pairs. Position represents the PCC between the expression distance of a spot to other spots and its physical distance to other spots. Random represents the PCC between the expression distance of a spot to other spots and a random spot physical distance to other spots. **E**. Schematic of image feature extraction. Fixed-size subimages for all spots are cropped according to their coordinates in the parallel H&E-stained image, and the image features, such as and, are extracted from these subimages. **F**. Heatmap shows that all spots in the same cluster tend to show similar image features. The image feature distances are normalized correlation-based distances of image features between all spot pairs. The colors are consistent with eleven clusters of spots in A. **G**. The plot shows the relationship between the expression distances and image feature distances for spot pairs. The expression distances are consistent with the description in C. The values in the plot are average expression distances s.d. **H**. The PCCs between expression distances and image feature distances for all spot pairs. The image feature represents the PCC between the expression distances of a spot to other spots and its image feature distances to other spots. Random represents the PCC between the expression distance of a spot to other spots and a random spot image feature distance to other spots. *** means p-value < 0.001.

### Workflow of ImSpiRE

Based on the above confirmed assumptions, we develop an image feature-aided spatial resolution enhancement method (ImSpiRE) for the ISC spatial transcriptome. ImSpiRE consists of three steps. In the first step, ImSpiRE crops the fixed-size subimages for all spots based on the spot coordinates of the parallel histology image. ImSpiRE then defines the patches, *i*.*e*., the virtual subspots, which are ranged densely on the parallel histology image. The equally sized subimages of patches are cropped in the same way as the spots. Then, for each spot or patch, a series of image features (such as intensity and texture, *etc*.) are extracted from its subimage. The size of the subimages determines the enhanced spatial resolution. For example, if the diameter of a spot in a histology image is approximately 200 pixels and the edge size of the subimages is set as 100 pixels, ImSpiRE can improve the resolution by approximately 3-4 times. In the second step, ImSpiRE computes the distance networks for the spots, the patches, and the spot-patch pairs. To build the spot network, ImSpiRE calculates an expression distance for each spot pair and then constructs a k-neighbors graph (see Methods for details). It then computes the shortest path lengths for each spot pair and finally builds a spot network (see Methods for details). To build the patch network, ImSpiRE uses a similar approach to obtain a physical distance network and an image feature distance network by using the coordinates and image features of all patches. ImSpiRE then combines both networks by a weight to build the patch network (see Methods for details). To build the spot-patch pair network, ImSpiRE uses a similar approach to construct two bipartite networks (*i*.*e*., a physical distance network and an image feature distance network) for all spot-patch pairs. Then, the same weight is used to combine two networks to obtain the spot-patch pair network. In the third step, ImSpiRE redistributes the gene expression from original spots to patches as follows. We formulate this task as the entropic regularized fused Gromov-Wasserstein transport (FGW) problem[17] and solve this optimal transport problem by using the networks in the second step, resulting in an optimal probabilistic embedding matrix that redistributes the expression profiles of each original spot to all patches (see Methods for details). Given the original gene expression profiles of spots and the derived probabilistic embedding matrix, ImSpiRE quantifies the gene expression of each patch as a weighted average of the original expression profile of all spots. ImSpiRE then normalizes the expression profiles of all patches to construct the transcriptional profiles with enhanced resolution (see Methods for details). ImSpiRE not only enhances the spatial resolution of the ISC spatial transcriptome to the subspot level but also imputes the expression profiles in unmeasured regions with the same resolution.

### Performance evaluation of ImSpiRE

To systematically evaluate the performance of ImSpiRE, we generate synthetic datasets. Briefly, we synthesize a large number of single-cell expression profiles with three cell types and a simulated parallel histology image with distinct image features for the three cell types (Fig. 3A, see Methods for details). We then assign single cells to coordinates in the image by cell type to simulate the tissue architecture and obtain a ground truth spatial transcriptome. To mimic the spots of the ISC spatial transcriptome, the gene expression levels of 9 nearby cells are merged as an observed spot, which means that the spatial resolution of the synthetic dataset is nine cells. We repeat the same procedure to generate 10 synthetic datasets and apply ImSpiRE to them to obtain transcriptome profiles with single-cell resolution. To quantitatively evaluate the performance of ImSpiRE, we group the obtained patches into three clusters, count the numbers of patches with different cell types to the majority in each cluster, and then calculate the F1 score and the adjusted Rand index (ARI) (see Methods for details). The results show that ImSpiRE effectively improves the resolution of synthetic datasets by nine times and accurately recovers the spatial information of three cell types (Fig. 3B, Fig. S2A). We further applied three widely used computational methods, CARD, BayesSpace and XFuse, to the same synthetic datasets and evaluated their performances, and ImSpiRE outperformed all of them in all cases (Fig. 3B, Fig. S2A).

**Fig. 2.**
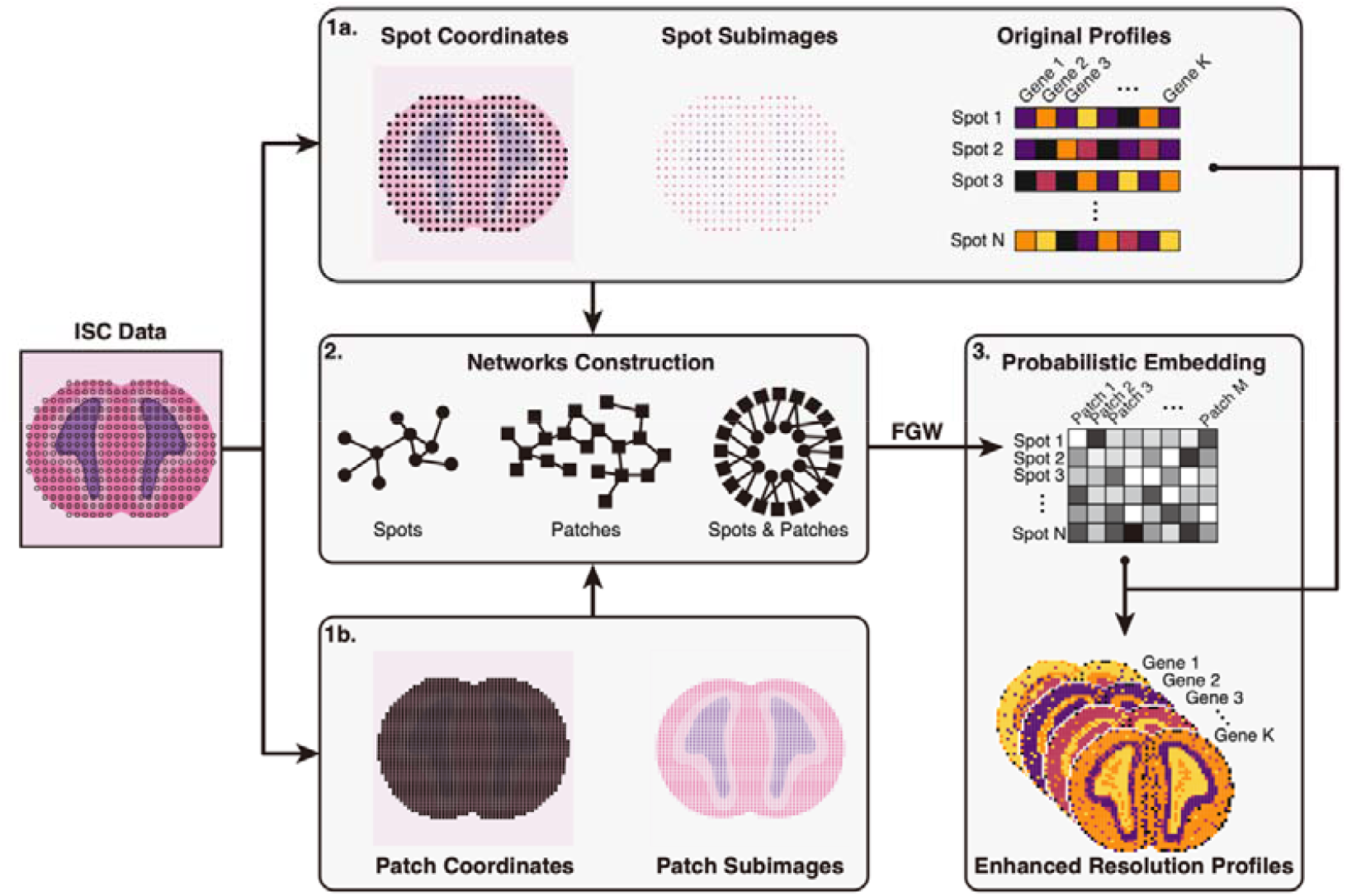
Workflow of ImSpiRE. ImSpiRE consists of three steps. In step 1a, ImSpiRE crops the fixed-size subimages for all spots based on the spot coordinates of the parallel histology image. In step 1b, ImSpiRE defines the patches, which range densely on the parallel histology image. The equally sized subimages of patches are cropped in the same way as the spots. For each spot or patch, a series of image features are extracted from its subimage. In step 2, ImSpiRE computes the distance networks of spots, patches, and spot-patch pairs by using the information derived in step 1. In step 3, ImSpiRE solves the entropic regularized fused Gromov-Wasserstein transport (FGW) problem to find an optimal probabilistic embedding matrix that redistributes the gene expression from original spots to patches. Given the original gene expression profiles of spots and the derived probabilistic embedding matrix, ImSpiRE quantifies the gene expression of each patch as a weighted average of the original expression profiles of all spots. ImSpiRE then normalizes the expression profiles of all patches to construct transcriptional profiles with enhanced resolution.

**Fig. 3.**
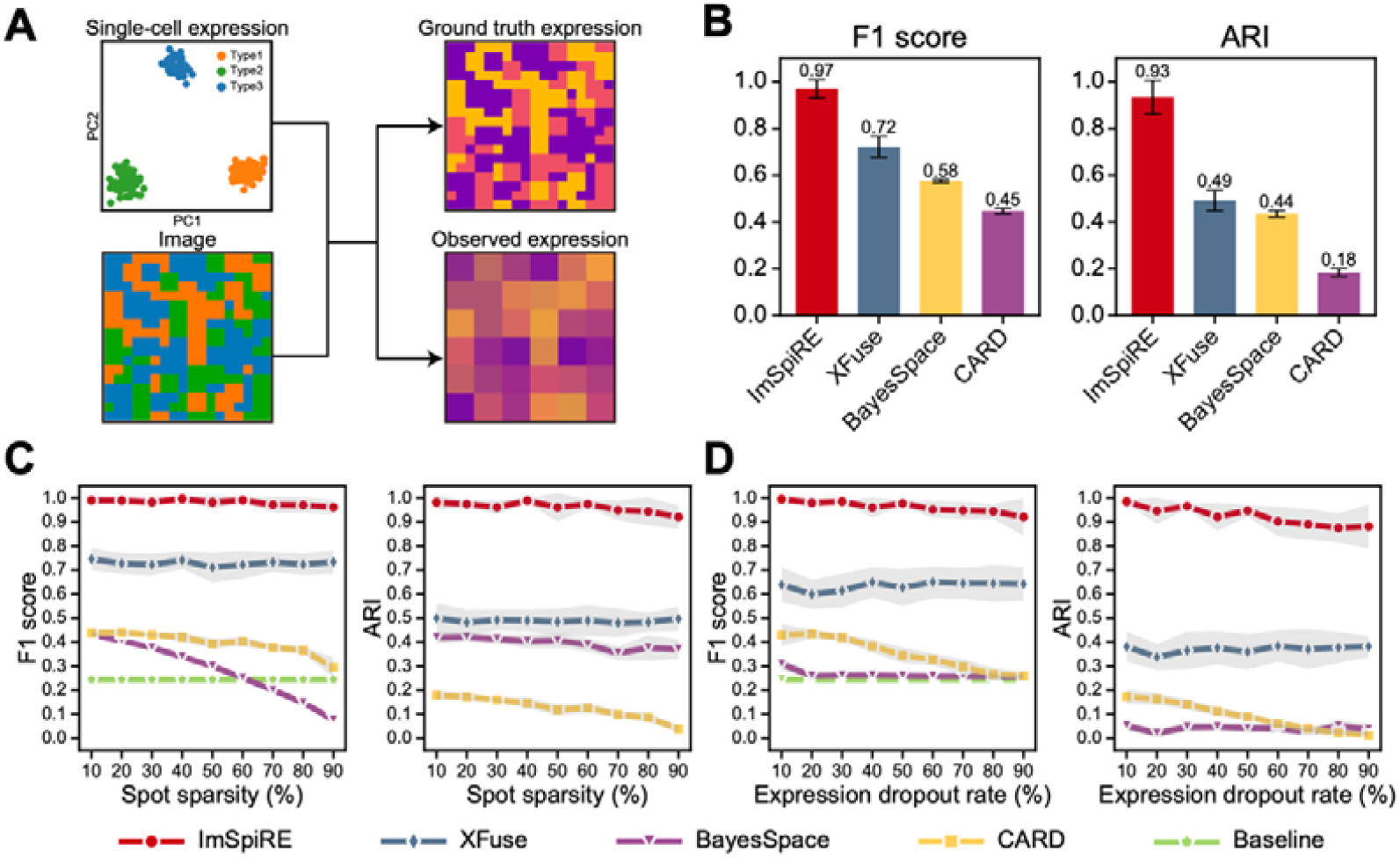
Performance evaluation of ImSpiRE. **A**. Design diagram of synthetic datasets. Left: a large number of single-cell expression profiles with three cell types and a simulated parallel histology image with distinct image features for the three cell types. Right: ground truth spatial transcriptome and observed spatial transcriptome. Single cells are assigned to coordinates in the image by cell type to obtain a ground truth spatial transcriptome, and the gene expression levels of each of the 9 nearby cells are merged as the observed spatial transcriptome. **B**. Performance of ImSpiRE and other methods on synthetic datasets. The values in bar plots are mean values and 95% confidence intervals for F1 score and ARI, respectively. **C**. Dynamic changes in F1 scores and ARIs in ImSpiRE and other methods with increasing spot sparsity. **D**. Dynamic changes in F1 scores and ARIs in ImSpiRE and other methods with increasing expression dropout. The lines of the plots in C and D are the mean values of the F1 scores and ARIs, respectively, and the gray shadows represent the 95% confidence intervals around the mean values. The baseline values are based on random prediction.

To evaluate the robustness of ImSpiRE for spot sparsity, we randomly remove a proportion of observed spots from the synthetic datasets and then apply ImSpiRE (see Methods for details). ImSpiRE shows consistently good performance even when only 10% of the spots remain (Fig. 3C, Fig. S2B). To evaluate the robustness of ImSpiRE for expression dropout, we randomly reset the expression values of a proportion of genes in synthetic datasets to 0. We apply ImSpiRE on these synthetic datasets, and its performance remains high, even when the dropout rate reaches 90% (Fig. 3D, Fig. S2C). We also compare the robustness of the other three methods with ImSpiRE, and ImSpiRE outperforms all the other methods on synthetic datasets (Fig. 3C, D, Fig. S2B, C).

### ImSpiRE enhances the spatial resolution to the subspot level

To further evaluate the spatial resolution enhancement performance on real datasets, we apply ImSpiRE to a widely used ST spatial transcriptome of the mouse olfactory bulb (MOB) [8], which contains 234 spots (Fig. 4A, B). In the parallel H&E-stained image of this dataset, 1 pixel represents approximately 0.68 μm. We set the edge size of the subimages as 50 pixels (*i*.*e*., ∼34 μm), which means that ImSpiRE can improve the spatial resolution by approximately 7 times (Fig. 4C). To evaluate the performance of ImSpiRE enhancement, *in situ* hybridization (ISH) data from the Allen Mouse Brain Atlas[18] are used as references (Fig. 4D, Fig. S3A). *Penk, Fabp7* and *Acsbg1* are expressed mainly in the granule cell layer, the nerve layer, and the mitral cell layer, respectively (Fig. 4D, Fig. S3A), but the original profiles are too coarse to obtain clear spatial patterns of these genes (Fig. 4E, Fig. S3A). ImSpiRE constructs transcriptional profiles with enhanced resolution, where those genes display consistent spatial patterns with ISH references (Fig 4F). Moran’s I test is applied in both the original and ImSpiRE-enhanced profiles to quantify the spatial autocorrelation for patterns of those genes (see Methods for details), and *Penk, Fabp7* and *Acsbg1* show much higher spatial autocorrelation in the ImSpiRE-enhanced profile than in the original profile (Fig. 4E, F, Fig. S3B). We further zoom in on a small portion of the dataset, and *Penk, Fabp7* and *Acsbg1* display subspot level spatial patterns within a spot in the ImSpiRE-enhanced profile (Fig. 4G), which is reasonably consistent with the fine structure of the tissue. Taken together, our results demonstrate that ImSpiRE can successfully enhance the ISC spatial resolution to the subspot level while refining the transcriptional landscapes.

**Fig. 4.**
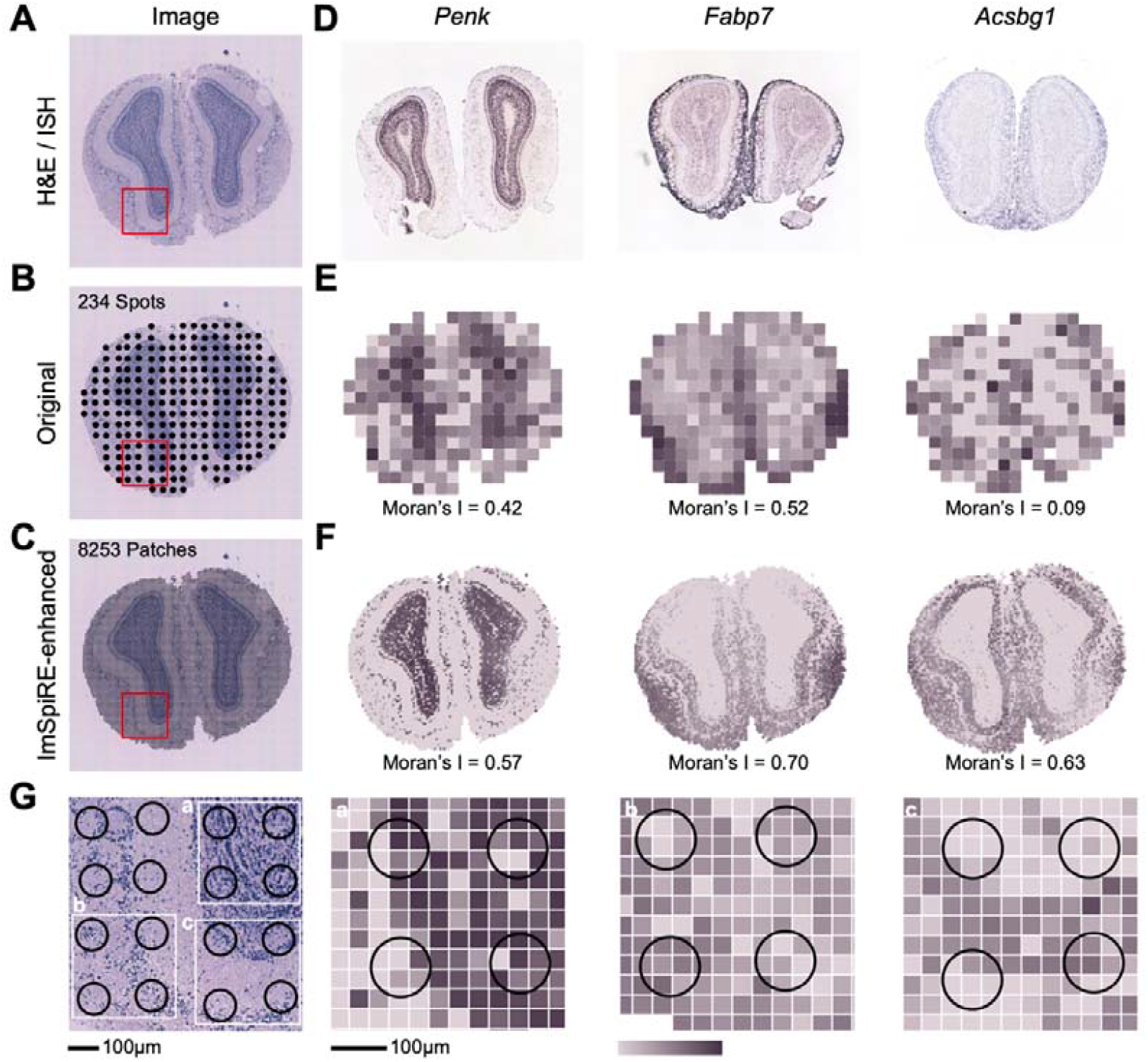
ImSpiRE enhances the spatial resolution to the subspot level in the MOB dataset. **A**. A parallel H&E-stained image of the MOB dataset. **B**. The coordinates of spots on the parallel H&E-stained image of the MOB dataset. **C**. The coordinates of patches on the parallel H&E-stained image of the MOB dataset. **D**. The reference ISH images of the MOB from the Allen Mouse Brain Atlas. **E**. The transcriptional profiles of spots with original resolution. **F**. The transcriptional profiles of patches with enhanced resolution. Spatial autocorrelations are listed at the bottom of the original and ImSpiRE-enhanced profiles in E and F. **G**. A zoomed parallel H&E-stained image of the red box with 16 spots. The black circles represent spots with a diameter of 100 μm. The three white boxes are small portions of the zoomed image. Approximately seven subimages can be observed within a single spot. The colors in E, F and G represent normalized values of gene expression.

### ImSpiRE benefits the identification of spatial domains and ligand□receptor interaction patterns

To evaluate the performance of ImSpiRE on tumor tissue with a complex spatial structure, we apply ImSpiRE to a Visium spatial transcriptome of squamous cell carcinoma (SCC) [19], which contains a tumor core, immune infiltrated stroma and mesenchyme stroma based on a manual annotation of its parallel H&E-stained image[11] (Fig. 5A). In its parallel H&E-stained image, 1 pixel represents approximately 0.28 μm. We set the edge size of the subimages as 70 pixels (*i*.*e*., ∼19.6 μm), which means that ImSpiRE can improve the spatial resolution by approximately 7 times. We separately cluster the original resolution spots and ImSpiRE-enhanced patches into 6 clusters according to their transcriptome similarity (Fig. 5A). According to the manual annotation, the clusters of C3 in both the original and ImSpiRE-enhanced profiles are related to the annotated tumor core. Compared with the original profile, cluster C1 in the ImSpiRE-enhanced profile clearly delineated the interface of the tumor core. The annotated immune infiltrated stroma is related well with cluster C2 in the ImSpiRE-enhanced profile, while it is not clearly related to any cluster in the original profile. To further characterize the clusters in the ImSpiRE-enhanced profile, we perform cluster-specific gene expression analysis and GO enrichment analysis (Fig. 5B, C). Consistent with the above description, clusters C1 and C2 are renamed the tumor core cluster (TCC) and immune infiltration cluster (IIC), respectively. Clusters C3-6 are related to the annotated mesenchyme stroma. As some angiogenesis-related genes (such as *COL18A1, COL4A2* and *COL4A1*) are highly expressed in cluster C2, we rename it the tumor adjacent cluster (TAC), and clusters 4-6 are renamed mesenchyme stroma clusters 1-3 (MSC1, MSC2 and MSC3). Our results demonstrate that ImSpiRE can benefit the identification of spatial domains in tissues with complex structures.

**Fig. 5.**
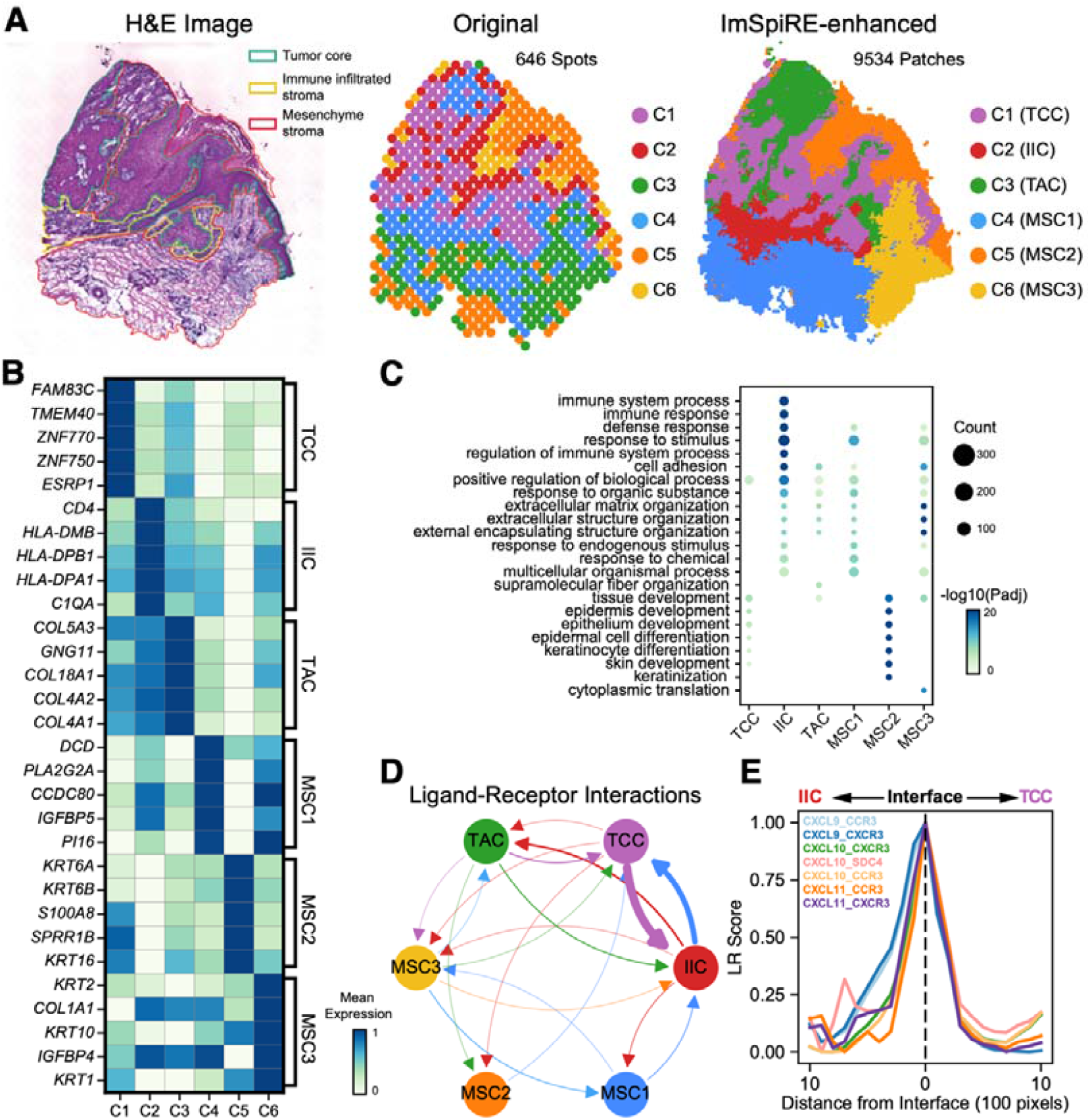
ImSpiRE allows the discovery of spatial domains and ligand□receptor interaction patterns in the SCC dataset. **A**. Left: Manual annotation of the parallel H&E-stained image of the SCC dataset. Three regions in this tissue are highlighted by different colors. Middle: Six clusters of spots at the original resolution. Right: Six clusters of patches at the enhanced resolution of ImSpiRE. **B**. Top 5 cluster-specific genes of patch clusters are shown in the heatmap. The values in the heatmap are the mean values of normalized gene expression. Clusters C1-C6 are defined as the tumor core cluster (TCC), immune infiltration cluster (IIC), tumor adjacent cluster (TAC) and mesenchyme stroma clusters (MSC1, MSC2 and MSC3), respectively. **C**. GO enrichment analysis in each cluster of patches. The size of the dot represents the number of genes enriched in the term, and the colors represent the enrichment significance. **D**. Ligand□receptor interaction network between 6 clusters in the SCC dataset. Arrows show the direction of interaction, and the linewidth represents the strength of the interaction. **E**. Ligand□receptor signaling communication at the interface of the TCC and IIC. The dashed line indicates the interface between the TCC and IIC. The lines of the plot are the mean values of the normalized LR scores.

As signaling communication occurs frequently in tumor tissues with complex structures[20], we further examine whether the ImSpiRE-enhanced profile can benefit the characterization of signaling communication. For the ImSpiRE-enhanced profile of the SCC dataset, we outline the ligand □ receptor interaction network among 6 clusters by using stLearn[21] (Fig. 5D, see Methods for details). There is a strong bidirectional link between TCC and IIC, which is reasonable because immune cells and tumor cells interact frequently in tumor tissues. With the improvement in interface resolution between TCC and IIC clusters, we further calculate the ligand □ receptor scores (LR scores) [21] at patches around the interface. We observe several distinct ligand □ receptor interaction spatial distribution patterns (Fig. 5E, Fig. S4). For example, CXCL9, -10, and -11/CXCR3 interactions occur exactly at the interface of the TCC and IIC (Fig. 5E), which is consistent with the role of CXCL9, -10, and -11/CXCR3 interactions in immune cell migration, differentiation, and activation[22]. Taken together, ImSpiRE not only aids in revealing complex spatial architectures but also benefits the characterization of the spatial distribution patterns of signaling communications.

### ImSpiRE improves spatiotemporal characterization for time-series spatial transcriptomes

Time-series spatial transcriptomes can provide both spatial and temporal information. To examine whether ImSpiRE can improve spatiotemporal characterization, we apply ImSpiRE to time-series ST spatial transcriptomes of human heart development (HHD)[23] containing three developmental stages (*i*.*e*., 4.5-5 postconception weeks (PCW), 6.5 PCW and 9 PCW). In the parallel H&E-stained images of this dataset, 1 pixel represents approximately 0.92 μm. We set the edge size of the subimages as 50 pixels (*i*.*e*., ∼46 μm), which means that ImSpiRE can improve the spatial resolution by approximately 4 times. To acquire the temporal order of spots or patches, we merge the profiles from three development stages and separately infer the pseudotime values for spots in the original profiles and patches in the ImSpiRE-enhanced profiles by Monocle3[24]. For the original profiles, the distributions of pseudotime values are indistinguishable among the three stages (Fig. 6A). In contrast, for ImSpiRE-enhanced profiles, the pseudotime values for patches in 4.5-5 PCW dataset tend to be much smaller than those of the other two datasets, while the values for patches in 9 PCW are much larger; this is consistent with the temporal order of development. The results demonstrate that spatial resolution enhancement of ImSpiRE can improve the characterization of temporal information for time-series spatial transcriptomes, probably due to the reduction in the mixing level of the original spots.

**Fig. 6.**
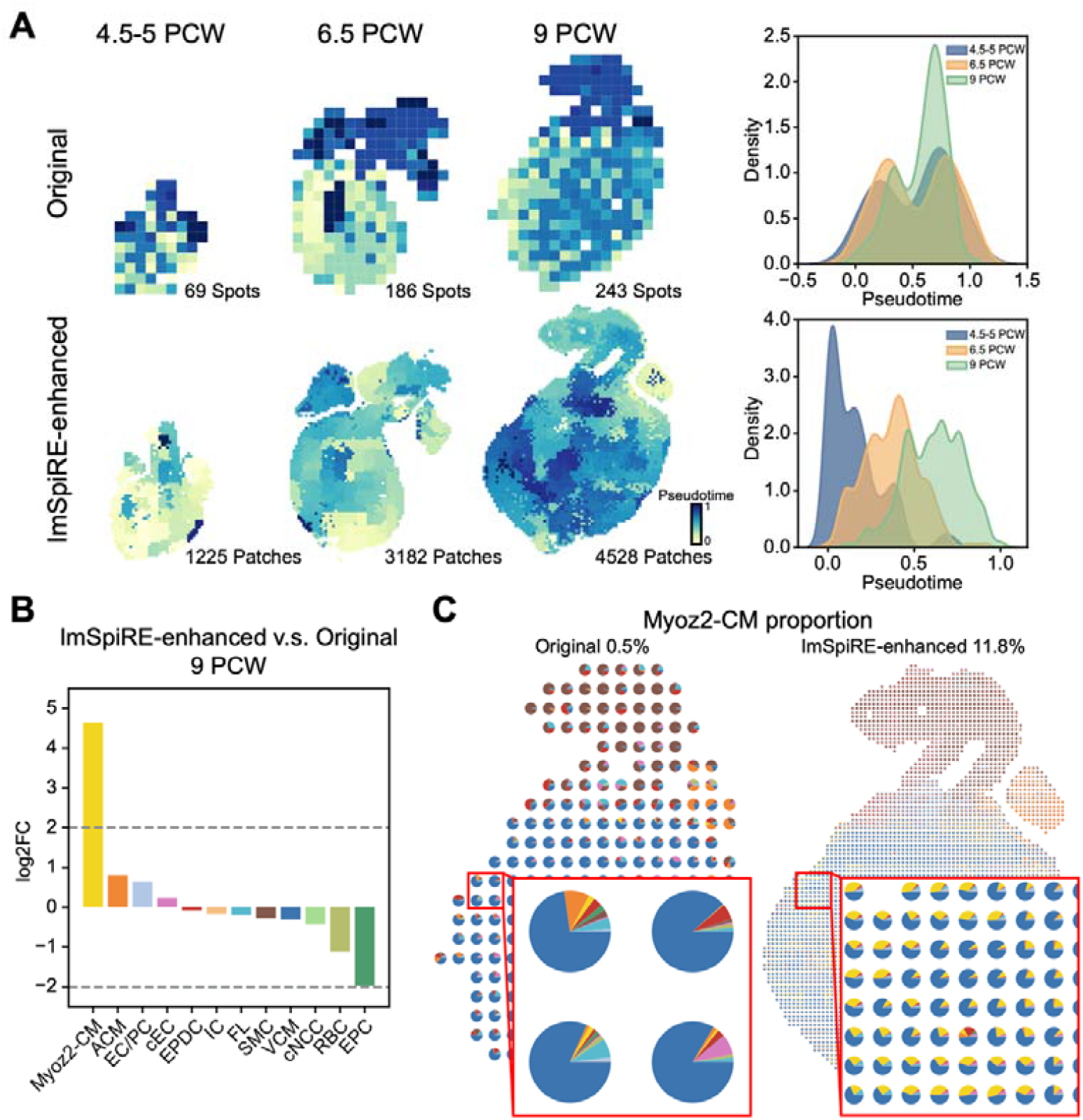
ImSpiRE improves the spatiotemporal characterization in HHD datasets. **A**. Top: Spatial distributions and density distributions of pseudotime at three development stages at the original resolution. Bottom: Spatial distributions and density distributions of pseudotime at the three stages at the enhanced resolution of ImSpiRE. The colors in spatial distributions represent the pseudotime, and three colors in density distributions of pseudotime represent three development stages. **B**. The proportions of cell types change between original and enhanced resolution at 9 PCW. The examined cell types included Myoz2-enriched cardiomyocytes (Myoz2-CM), atrial cardiomyocytes (ACM), endothelium/pericytes (EC/PC), capillary endothelium (cEC), epicardium-derived cells (EPDC), immune cells (IC), fibroblast-like (FL), smooth muscle cells (SMC), ventricular cardiomyocytes (VCM), cardiac neural crest cells (cNCC), red blood cells (RBC) and epicardial cells (EPC). Dashed lines indicate the presence of 4-fold changes. **C**. Spatial distribution of Myoz2-CM proportion between original and enhanced resolution at 9 PCW. Pie charts represent cell type proportions in spots or patches. The colors in pie charts represent different cell type proportions that correspond to the colors in B.

As the resolutions of the parallel H&E-stained images are relatively low (∼0.92 μm/pixel) for the HHD datasets, ImSpiRE-enhanced profiles cannot achieve single-cell resolution. We suspect that such a nonsingle-cell-resolution enhancement can benefit the performance of cell-type deconvolution. To assess the above assumption, we applied CARD[10] for cell-type deconvolution on the original and ImSpiRE-enhanced profiles at 9 PCW (see Methods for details). Comparing the proportions of cell types between the original and ImSpiRE-enhanced profiles, we observe that Myoz2-enriched cardiomyocytes (Myoz2-CM) are abundantly detected in the ImSpiRE-enhanced profile (0.5% in the original profile *vs*. 11.8% in the ImSpiRE-enhanced profile; Fig. 6B), which is consistent with a recent estimation that the population of Myoz2-CM is over 10% in the human embryonic heart[23]. When zooming in on a ventricle region, we observe high proportions of Myoz2-CM in ImSpiRE-enhanced patches but very low proportions in original spots (Fig. 6C), probably because the lower mixing level in the patches can increase the sensitivity of distinguishing similar cell types. Taken together, our results demonstrate that ImSpiRE is advantageous for accurate cell type identification in the presence of prior knowledge.

## Discussion

In this study, we developed ImSpiRE, an image feature-aided spatial resolution enhancement method, for the ISC spatial transcriptome. Without any additional data except a parallel histology image, ImSpiRE constructs the subspot resolution transcriptional profiles while imputing gene expression of the unmeasured tissue regions. To the best of our knowledge, most, if not all, ISC spatial transcriptomes contain parallel histology images of tissue sections, which guarantees the applicability of ImSpiRE. The parallel histology images (usually H&E-stained images) provide valuable information, such as cell morphology, staining intensity and texture. ImSpiRE leverages those image-derived features to enhance resolution. Compared to deep learning models, ImSpiRE ensures good interpretability of the results. The pixel density of the parallel histology image determines the upper limit of the resolution enhancement of ImSpiRE, as enough pixels are required for image feature extraction in subimages. For spatial transcriptomes with high-pixel-density parallel histology images, ImSpiRE-enhanced profiles could theoretically achieve single-cell resolution.

In the network construction step of the ImSpiRE workflow, information from both the physical distance network and image feature distance network is combined through a weighting parameter with default value 0.5. For certain tissues with complex architecture, in which the arrangement of different cell types is complicated, we recommend increasing the value of the weighting parameter to take advantage of the parallel histology images. In contrast, for special cases with indefinite information derived from parallel histology images (for example, due to low-quality H&E staining), the users can decrease the value of the weighting parameter to avoid the redistribution of gene expression of spots to irrelevant patches.

The usage of ImSpiRE can be easily combined with other single-cell transcriptome or spatial transcriptome analysis methods. For example, as we show in this study, applying pseudotime analysis, cell-type deconvolution analysis and signaling communication analysis to ImSpiRE-enhanced profiles can improve the inference of temporal order, the identification of cell types and the characterization of signaling communications. Other methods, such as trajectory inference and RNA velocity, can also be easily applied to ImSpiRE-enhanced profiles. In addition, the ImSpiRE-enhanced profile can be combined with other types of images, such as spatial proteomic[25] or spatial metabolomic images[26], to provide insights into the correlation between spatial architecture and complex cellular behavior.

## Methods

### Data preprocessing

For all public datasets, we obtained the raw gene expression matrices, spot coordinates, clustering results and parallel histology images from the 10x Genomics data repository or from publications (see Supplementary Information for details). For the ST datasets of human heart development, we converted the Ensemble IDs to gene names and filtered out the genes that do not match. Then, we averaged the expression values for the duplicated genes. For all original gene expression profiles, we retained the genes that were expressed in at least 10 spots and took the spots that were located in the tissue section. Details of dataset availability are provided in the Supplementary Information.

### Assumption validation

For each dataset used in the validation, we performed principal component analysis (PCA) on the gene expression profiles and selected the top 15 PCs to calculate the pairwise correlation-based distance of gene expression, which is called the expression distance. The Euclidean distance, *i*.*e*., physical distance, was calculated according to the spot coordinates. We then cropped the fixed-size subimages with an edge size of 200 pixels for all spots based on the spot coordinates of the parallel histology image. We used CellProfiler[27], a cell image analysis software, to extract features for all the subimages. We also performed PCA on the image features and selected the top 15 PCs to calculate the pairwise correlation-based distance, which is called the image feature distance. All distances were scaled to be between 0 and 1. We calculated the Pearson correlation coefficients (PCCs) between the expression distance of each spot to other spots and its physical distances to other spots. To verify the significance, we calculated the PCCs between the expression distances of each spot to other spots and a random spot’s physical distances to other spots. A similar approach was applied to calculate the correlation between expression distance and image feature distance. All datasets were validated as described above.

### ImSpiRE workflow

ImSpiRE consists of three steps to construct the enhanced resolution profiles. In the first step, ImSpiRE crops the fixed-size subimages for all *N* spots based on these spot coordinates of the parallel histology image. Then, ImSpiRE densely arranges *M* patches at intervals equal to the edge size of subimages on the parallel histology image and crops equally sized subimages of patches. ImSpiRE provides two image feature extraction modes. The first mode directly extracts the intensity and texture features of the image, and the second mode extracts image features using CellProfiler with the preset pipelines. For each spot or patch, a series of image features are extracted from its subimage, retaining only those with a standard deviation > 1. In the second step, ImSpiRE computes the distance networks of spots, patches, and spot-patch pairs.To build the spot network, ImSpiRE performs PCA on the original resolution expression profiles with *N* spots and *g* genes, represented by a matrix *P* ∈*R*^*N* × *g*^, and selects the top 30 PCs to calculate the pairwise correlation-based distance between all spots. ImSpiRE uses this pairwise distance to construct a k-neighbors graph and computes the shortest path lengths from this graph for each pair of spots, resulting in a graph-based expression distance network, *G*^Spot^ ∈*R*^*N*×*N*^ (*i*.*e*., the spot network). To build the patch network, we choose Euclidean distance and correlation-based distance as the distance metric for physical distance and image feature distance, respectively, and use a similar approach to construct the graph-based physical distance network *G*^phy^ ∈*R*^*M*×*M*^ and the graph-based image feature distance network *G*^img^ ∈*R*^*M*×*M*^ for patches. To obtain the patch network, we consider the following:

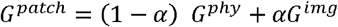

where α ∈ [0,1] is a weighting parameter to balance the weight between *G*^phy^ and *G*^img^. To build the spot-patch pair network, ImSpiRE uses the same approach to construct two bipartite networks (*i*.*e*., a graph-based physical distance network and a graph-based image feature distance network) for all spot-patch pairs, and α is used to combine two networks to obtain the spot-patch pair network *G*^Spot patch^ ∈*R*^*N*×*M*^. In the third step, we find a probabilistic embedding matrix 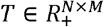 to redistribute the expression profiles of each original spot to all patches. We formulate this task as a generalized optimal transport problem[28, 29], and the optimization requirements over T are as follows.

To ensure that nearby patches with similar image features can share similar levels of gene expression after redistributing, we measure the pairwise discrepancy of *T* for the spot network *G*^Spot^and the patch network *G*^patch^ by using the Gromov-Wasserstein discrepancy[30]

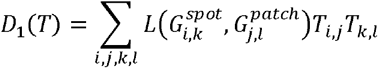

Where *L* is a quadratic loss function, *i, k* are spots, and *j, l* are patches. For each spot *i*, the value of *T*_*i, j*_ is the relative probability of embedding it to patch *j*. This term shows the preference to embed spots such that if spots *i* and *k* are embedded into patches *i* and *j*, respectively, the expression distance between *i* and *k* (*i*.*e*., 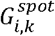) corresponds to the physical distance and image feature distance between *j* and *l* (*i*.*e*., 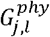 and 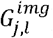). To ensure that each spot can be redistributed to patches that are close to it and to have consistent image features, we consider

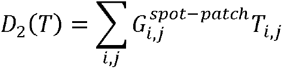

This term represents the discrepancy of *T* for *G*^Spot–patch^. Intuitively, if a spot *i* is embedded into a patch *j*, then *j* should be close to *i*,, and similar subimages should be observed between *i* and *j*.

Furthermore, a spot cannot be embedded into only a few patches because of the number difference between N spots and M patches. We regularize *T* by favoring embeddings with higher entropy to ensure that all patches are embedded properly. An entropic regularized term[29] is defined as

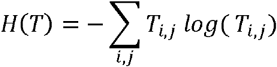

Putting these together, we define the entropic regularized fused Gromov-Wasserstein transport (FGW) problem[17] for the optimal probabilistic embedding *T*^*^:

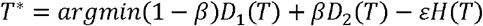

where β ∈ [0,1] is a constant interpolating parameter and *ε* is a nonnegative regularization constant. *T* should be consistent with the marginal distributions over the spots and patches, *s* and *p*, respectively. In this case, and can be set to be uniform distributions. It is subject to

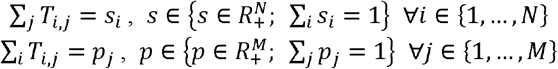

The optimal probabilistic embedding *T*^*^ minimizes (1 − β)*D*_1_ (*T*) + β D_2_ (*T*) and simultaneously maximizes *εH*(*T*). We use an open-source Python package POT[31] to calculate *T*^*^. Given the original resolution expression profiles *P*∈*R*^*N*×*g*^, we can derive expression profiles, *P*′ = *P*^*T*^*T*^*^ ∈ *R*^*g*×*M*^, which contain the relative expression values of g genes for M patches. We then normalize the expression profiles for each patch to construct the transcriptional profiles with enhanced resolution. For example, if 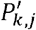 represents the relative expression value for gene *k* in patch *j*, we normalize it as

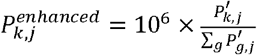

### Synthetic datasets

We used the R package Splatter to simulate single-cell datasets containing 10,000 cells of three cell types with 2,000 genes. We then use Splatter to generate another dataset corresponding to four different types, including three cell types and background. We perform PCA on this dataset, and the top 15 PCs are selected as the image features of the three cell types and background. We assign these single cells and image features at 3,600 locations (a matrix of 60×60 cells) to obtain a ground truth spatial transcriptome and a simulated parallel histology image. Then, we sum the gene expression values of 9 adjacent cells (a matrix of 3×3 cells) to obtain 400 spots (a matrix of 20×20 spots) as the observed spatial transcriptome. All 10 synthetic datasets are generated as described above.

### Method evaluation

We compare three other methods with ImSpiRE on synthetic datasets, including CARD (v1.0), BayesSpace (v1.4.1) and XFuse (v0.2.1). We take the transcriptome profiles of all spots as the input and use the four methods to increase the spatial resolution to obtain a 60×60 matrix, corresponding to the ground truth spatial transcriptome. The input spots with all gene expression values equal to 0 are removed. We provide additional input for these methods if needed. For CARD, we input the single-cell data from the ground truth spatial transcriptome. For XFuse, we input the image of cell types recorded in different colors. To quantitatively evaluate the performance of the methods, we use the scanpy.tl.louvain() in the Python package SCANPY[32] to perform Louvain clustering analysis on the transcriptional profiles constructed by ImSpiRE, CARD and XFuse. For BayesSpace, we use its own clustering results. All constructed profiles are appropriately classified into 3 clusters and compared with the ground truth cell types to calculate the F1 scores and ARIs.

To evaluate the robustness for spot sparsity, we randomly remove 10%-90% of the observed spots from the synthetic datasets while preserving all other additional information and then apply the four methods. Missing spots are counted as incorrect predictions. To evaluate the robustness for gene expression dropout, we randomly set the expression values of 10%-90% of the genes that are not 0 to 0 in each spot of the synthetic datasets and retain other additional information. Then, F1 scores and ARIs are calculated using the same approach.

### Gene spatial autocorrelation test

To quantify the spatial autocorrelation for patterns of genes, we first select the top 1,000 highly variable genes in the original resolution profile of the MOB dataset. Then, we use geopandas. GeoDataFrame() in the Python package geopandas[33] to convert the spot coordinates to GeoDataFrame and use esda.moran.Moran() in the Python package esda[34] to calculate Moran’s I of these genes in both the original resolution profile and the ImSpiRE-enhanced profile. We further select genes based on the p-value and z-score of Moran’s I and keep the genes with p-value < 0.05 and |z-score| > 1.96.

### Cluster-specific gene analysis and GO enrichment analysis

In this analysis, we find the cluster-specific genes of 6 clusters by using scanpy.tl.rank_genes_groups() in the Python package SCANPY. We apply g:Profiler[35] to enrich the biological processes for 6 clusters by using their top 200 cluster-specific genes.

### Ligand□receptor interaction analysis

In this analysis, stLearn[21], a downstream toolkit for spatial transcriptomics data, is used to infer the interaction network. Databases containing 2,293 ligand □ receptor candidates are available within stLearn. Of these, 1,659 ligand □ receptor pairs are found in the SCC dataset. For the ImSpiRE-enhanced resolution profile, we keep the genes that are expressed in at least 3 patches, and we use stlearn.pp.normalize_total() to normalize the total count of each patch to the median of total counts for patches before normalization. Then, we use st.tl.cci.run_cci() to perform the ligand □ receptor analysis provided in the stLearn package to calculate LR scores for each patch in 1,659 ligand □ receptor pairs. We calculate the minimum Euclidean distance of the pixels in the image between all IC patches and TC patches as the distance of the patches from the cluster edge. We divide these patches into 21 bins based on the pixel distance away from the edge. The LR scores of the patches in each bin are averaged as the LR score in that pixel distance bin.

### Pseudotime analysis

We merge the original resolution profiles of the three stages of human heart development together and remove the batch effect using align_cds()[36] in the R package monocle3. When inferring the pseudotime, a spot or a patch in the 4.5-5 PCW stage is randomly chosen as the start. We scale the pseudotime of all spots or patches in the three stages to be 0-1 for comparison between the original resolution profiles and the ImSpiRE-enhanced profiles.

### Cell type deconvolution

We use the human embryonic heart scRNA-seq data from [23] as a reference and CARD to deconvolve cell types. For both the original resolution profile and the ImSpiRE-enhanced profile at the 9 PCW stage, we first keep genes that are expressed in at least 10 spots or patches and create a CARD object by using createCARDObject(). Then, we use CARD_deconvolution() to infer the proportion of cell types for each spot or patch. According to previous studies, 12 different cell types have been defined in scRNA-seq data. We count the average proportion of these 12 cell types across all spots or patches as the total cell type proportion for the 9 PCW stage.

## Data availability

All datasets analyzed in this study are publicly available (see Supplementary Information for details).

## Code availability

ImSpiRE is available as a Docker image at https://hub.docker.com/r/tongjizhanglab/imspire, and the source code is publicly available at https://github.com/TongjiZhanglab/ImSpiRE.

## Author contributions

Y(Yong).Z. conceived the project; Y.H. and Y(Yizhi).Z. performed method development and data analysis with the help of Z.G. and S.B; Y.H., Y(Yizhi).Z. and Y(Yong).Z. wrote the manuscript.

## Acknowledgements

This work was supported by the National Key Research and Development Program of China (2021YFA1302500) and the National Natural Science Foundation of China (32030022, 31970642).

## References

1. Lubeck, E., et al., Single-cell in situ RNA profiling by sequential hybridization. Nat Methods, 2014. 11(4): p. 360–1.

2. Shah, S., et al., In Situ Transcription Profiling of Single Cells Reveals Spatial Organization of Cells in the Mouse Hippocampus. Neuron, 2016. 92(2): p. 342–357.

3. Eng, C.L., et al., Transcriptome-scale super-resolved imaging in tissues by RNA seqFISH. Nature, 2019. 568(7751): p. 235–239.

4. Chen, K.H., et al., RNA imaging. Spatially resolved, highly multiplexed RNA profiling in single cells. Science, 2015. 348(6233): p. aaa6090.

5. Zhao, E., et al., Spatial transcriptomics at subspot resolution with BayesSpace. Nat Biotechnol, 2021. 39(11): p. 1375–1384.

6. Moses, L. and L. Pachter, Museum of spatial transcriptomics. Nat Methods, 2022. 19(5): p. 534–546.

7. Asp, M., J. Bergenstrahle, and J. Lundeberg, Spatially Resolved Transcriptomes-Next Generation Tools for Tissue Exploration. Bioessays, 2020. 42(10): p. e1900221.

8. Stahl, P.L., et al., Visualization and analysis of gene expression in tissue sections by spatial transcriptomics. Science, 2016. 353(6294): p. 78–82.

9. Chen, A., et al., Spatiotemporal transcriptomic atlas of mouse organogenesis using DNA nanoball-patterned arrays. Cell, 2022. 185(10): p. 1777–1792 e21.

10. Ma, Y. and X. Zhou, Spatially informed cell-type deconvolution for spatial transcriptomics. Nat Biotechnol, 2022. 40(9): p. 1349–1359.

11. Sun, D., et al., STRIDE: accurately decomposing and integrating spatial transcriptomics using single-cell RNA sequencing. Nucleic Acids Res, 2022. 50(7): p. e42.

12. Cable, D.M., et al., Robust decomposition of cell type mixtures in spatial transcriptomics. Nat Biotechnol, 2022. 40(4): p. 517–526.

13. Elosua-Bayes, M., et al., SPOTlight: seeded NMF regression to deconvolute spatial transcriptomics spots with single-cell transcriptomes. Nucleic Acids Res, 2021. 49(9): p. e50.

14. Bergenstrahle, L., et al., Super-resolved spatial transcriptomics by deep data fusion. Nat Biotechnol, 2022. 40(4): p. 476–479.

15. Monjo, T., et al., Efficient prediction of a spatial transcriptomics profile better characterizes breast cancer tissue sections without costly experimentation. Sci Rep, 2022. 12(1): p. 4133.

16. Nitzan, M., et al., Gene expression cartography. Nature, 2019. 576(7785): p. 132–137.

17. Vayer, T., et al., Optimal Transport for structured data with application on graphs. International Conference on Machine Learning, Vol 97, 2019. 97.

18. Lein, E.S., et al., Genome-wide atlas of gene expression in the adult mouse brain. Nature, 2007. 445(7124): p. 168–76.

19. Ji, A.L., et al., Multimodal Analysis of Composition and Spatial Architecture in Human Squamous Cell Carcinoma. Cell, 2020. 182(2): p. 497–514 e22.

20. He, G., et al., Exosomes in the hypoxic TME: from release, uptake and biofunctions to clinical applications. Mol Cancer, 2022. 21(1): p. 19.

21. Pham, D., et al., stLearn: integrating spatial location, tissue morphology and gene expression to find cell types, cell-cell interactions and spatial trajectories within undissociated tissues. biorxiv, 2020.

22. Tokunaga, R., et al., CXCL9, CXCL10, CXCL11/CXCR3 axis for immune activation - A target for novel cancer therapy. Cancer Treat Rev, 2018. 63: p. 40–47.

23. Asp, M., et al., A Spatiotemporal Organ-Wide Gene Expression and Cell Atlas of the Developing Human Heart. Cell, 2019. 179(7): p. 1647–1660 e19.

24. Qiu, X., et al., Reversed graph embedding resolves complex single-cell trajectories. Nat Methods, 2017. 14(10): p. 979–982.

25. Lundberg, E. and G.H.H. Borner, Spatial proteomics: a powerful discovery tool for cell biology. Nat Rev Mol Cell Biol, 2019. 20(5): p. 285–302.

26. Yuan, Z.Y., et al., SEAM is a spatial single nuclear metabolomics method for dissecting tissue microenvironment. Nature Methods, 2021. 18(10): p. 1223-+.

27. Stirling, D.R., et al., CellProfiler 4: improvements in speed, utility and usability. BMC Bioinformatics, 2021. 22(1): p. 433.

28. Villani, C., Topics in optimal transportation. Vol. 58. 2021: American Mathematical Soc.

29. Villani, C., Optimal transport: old and new. Vol. 338. 2009: Springer.

30. Mémoli, F., On the use of Gromov-Hausdorff distances for shape comparison. 2007.

31. Flamary, R., et al., POT: Python Optimal Transport. J. Mach. Learn. Res., 2021. 22(78): p. 1–8.

32. Wolf, F.A., P. Angerer, and F.J. Theis, SCANPY: large-scale single-cell gene expression data analysis. Genome Biol, 2018. 19(1): p. 15.

33. Jordahl, K., et al., geopandas/geopandas: v0. 8.1. Zenodo, 2020.

34. Rey, S.J. and L. Anselin, PySAL: A Python library of spatial analytical methods, in Handbook of applied spatial analysis. 2010, Springer. p. 175–193.

35. Raudvere, U., et al., g:Profiler: a web server for functional enrichment analysis and conversions of gene lists (2019 update). Nucleic Acids Research, 2019. 47(W1): p. W191–W198.

36. Haghverdi, L., et al., Batch effects in single-cell RNA-sequencing data are corrected by matching mutual nearest neighbors. Nat Biotechnol, 2018. 36(5): p. 421–427.

